# Development and structure-guided characterization of a novel ACE2-binding macrocyclic peptide

**DOI:** 10.64898/2025.12.01.690145

**Authors:** Roger M. Benoit, Jinling Wang, Darja Beyer, Ata Abbas, Matthew J. Rodrigues, Mara M. Wieser, Xavier Deupi, Cristina Müller, Hiroaki Suga, Jeffrey W. Bode

## Abstract

Angiotensin-converting enzyme 2 (ACE2) is a key node in the protective axis of the renin-angiotensin-aldosterone system (RAAS) for blood pressure and hydroelectrolyte regulation, and the main protein receptor recognized by the spike glycoproteins of the severe acute respiratory syndrome (SARS) coronaviruses (CoV) SARS-CoV and SARS-CoV-2. We identified the macrocyclic peptide WJL-63, developed using mRNA display, with high ACE2-binding affinity. The peptide was characterized *in vitro* in terms of purity, stability, hydrophilicity and ACE2 binding. The crystal structure of the extracellular region of ACE2 in complex with the peptide at 2.2 Å resolution was elucidated. The structure revealed a binding mode in which WJL-63 is accommodated towards one side of the wide catalytic cleft of the ACE2 peptidase domain, with no direct contact to the conserved zinc ion site. WJL-63 residues Q4, R7, R11 and R14 anchor the peptide deep inside the binding pocket. The opposite edges of the peptide were found to be in contact with subdomain 1 and subdomain 2 of the peptidase domain. This upright binding mode requires an open ACE2 conformation, in contrast to small molecule carboxypeptidase inhibitors, which typically bind to the closed conformation of the enzyme. As a consequence of the open conformation binding mode, the front edge of WJL-63 is accessible for modification such as the herein reported conjugation of a chelator for radiometal labeling. The radiolabeled DOTA-WJL-63 was evaluated on ACE2-transfected HEK cells on which it revealed relatively strong binding with a K_D_ value of 90 ± 28 nM. WJL-63 provides a strong basis for the development of new classes of compounds for the modulation of ACE2 conformation, and for the development of imaging agents for the visualization of ACE2, for example in fluorescence or electron microscopy, or positron emission tomography (PET) imaging.

## Introduction

Our vascular system is closed and pressurized while the blood volume and the systemic vascular resistance are tightly regulated through complex mechanisms.^1^ The renin-angiotensin-aldosterone system (RAAS) is a key node for the regulation of blood pressure and systemic vascular resistance. Angiotensin-converting enzyme (ACE) and angiotensin-converting enzyme 2 (ACE2) are two major players in the RAAS system:^2^ ACE converts the decapeptide angiotensin I into angiotensin II which contains 8 amino acid residues. The latter binds to the angiotensin II type 1 receptor (AT_1_R), resulting in vasoconstriction and an increase in blood pressure. ACE2 removes another amino acid from angiotensin II, producing angiotensin (1-7). This peptide binds to the G-protein-coupled receptor (GPCR) proto-oncogene Mas, resulting in vasodilation and decreased blood pressure, along with anti-inflammatory and tissue-protective effects.^3, 4^ The ACE2 axis can, therefore, be viewed as a protective node of the RAAS system.^5^ Screening of peptides to identify ACE2 hydrolytic activity revealed that the enzyme can remove the C-terminal residue of at least eleven biological peptides, including the efficient hydrolyzation of angiotensin II, apelin-13, dynorphin A 1-13 and des-Arg^9^-bradykinin.^6^ A pH of 6.5 is optimal for proteolytic activity, and there is increased activity of ACE2 in the presence of Cl^-^and F^-^ anions.^6^

To date, only a few selective ACE2 binders have been identified, whereas a wide range of molecules are known with inhibitory activity for enzymes within the RAAS, most notably ACE.^7, 8^ This is partially due to the late discovery of ACE2, which occurred more than 40 years after the identification of ACE,^9–11^ but also due to the high similarity between these two enzymes which rendered the development of ACE2-selective ligands challenging. Initially, ACE2 inhibitors were developed with the intention to obtain a new class of therapeutic agents for the treatment of hypertension, similar to the well-known ACE inhibitors.^8^ However, based on the opposite physiological effects of ACE2 and ACE, ACE2 inhibitors would rather present a risk to cause hypertensive episodes, hence, this class of compounds was not further developed.^12^ Three main classes of ACE2 inhibitors are known: phosphinic peptide-based inhibitors,^13^ thiol-based inhibitors^14^ and non-peptidic small molecule inhibitors, such as MLN-4760.^15^ In 2003, the ACE2-selective cyclic peptide DX600 was developed by phage display technology to further investigate the function of ACE2 *in vivo*.^16^ Very recently, bicyclic peptides have been described that bind within the ACE2 cleft.^17^ Cyclic peptides can be particularly advantageous in view of their biological properties because of their structural rigidity leading to higher affinity and target selectivity as well as an improved stability compared to their linear analogues.^18^

With the aim to develop novel ACE2 radiotracers, we utilized the “Random non-standard Peptides Integrated Discovery” (RaPID) platform^19^ to generate a macrocyclic peptide, WJL-63, that specifically binds to ACE2. The complex formed between the extracellular segment of ACE2 and WJL-63 was biochemically, biophysically and structurally characterized.

## Results

### RaPID display campaign for peptide selection

We employed the RaPID system, which enabled the display of libraries of over 10^12^ unique macrocyclic peptides^20–22^ to identify candidate peptides based on their ability to bind tightly to ACE2. A puromycin-ligated mRNA library was constructed to encode peptides with N-chloroacetyl-D-tyrosine as the initiator amino acid followed by a random peptide region consisting of 6–15 residues followed by cysteine and ending with a short linker peptide. Upon translation of these mRNAs using the flexible *in vitro* translation (FIT) system, the chloroacetyl group on the N-terminus of the linear peptides undergoes a spontaneous reaction with the thiol group on the downstream cysteine to form thioether-macrocyclic peptides. Each macrocyclic peptide in the library is covalently linked to its corresponding mRNA via the puromycin linker for later amplification and identification by sequencing. These libraries were screened against His-tagged ACE2 immobilized on suitable magnetic Dynabeads.

After six iterative rounds of negative and positive selections, the recovery rate of peptide–mRNA fusion molecules from positive selections was sufficiently increased, suggesting that the population of cyclic peptides binding with ACE2 were selectively enriched. Next Generation Sequencing (NGS) analysis of the enriched libraries showed unique families of sequences, and we selected WJL-63 as our primary choice which had the highest frequency of appearance in the sequencing data.

The peptide WJL-63 was synthesized using solid-phase peptide synthesis (SPPS) and obtained with a purity greater than 99%, as confirmed by high-performance liquid chromatography (HPLC). The molecular mass was verified by HRMS (ESI) ([M+H]^2+^: m/z_calc_ = 1235.0582; m/z_found_ = 1235.0579). Cyclization of the linear peptide was carried out using tris(2-carboxyethyl)phosphine (TCEP) and triethylamine, followed by HPLC purification to achieve >99% purity. The cyclic peptide was confirmed by HRMS (ESI) ([M+H]^2+^: m/z_calc_ = 1217.0698; m/z_found_ = 1217.0723). The purified cyclic peptide was subsequently used for crystallization studies. The peptide DOTA–WJL-63 was synthesized by manual conjugation of the chelator DOTA-tris(tert-butyl ester) to the lysine side chain of WJL-63. HPLC analysis indicated a purity exceeding 99%. Its molecular mass was confirmed by HRMS (ESI) ([M+H]^4+^: m/z_calc_ = 705.5836; m/z_found_ = 705.5844).

### Radiolabeling and *in vitro* characterization of [^67^Ga]Ga-DOTA-WJL-63

[^67^Ga]Ga-DOTA-WJL-63 was obtained with high radiochemical purity of >96% at a molar activity up to 20 MBq/nmol. The radiopeptide was radiolytically stable over a period of 24 h (93 ± 5% intact radiopeptide). The addition of ascorbic acid to the radiopeptide formulation as a radical scavenger prevented this radiolytic degradation entirely resulting in >99% intact radiopeptide even after 24 h incubation. In mouse blood plasma, the radiopeptide was degraded, resulting in only 91% and 69% intact radiopeptide after 1 h and 3 h, respectively. The radiopeptide remained, however, completely stable in human blood plasma for to the entire period of investigation over 24 h (Figure 1A). A logD value of −4.4 ± 0.1 was determined for [^67^Ga]Ga-DOTA-WJL-63 indicating a hydrophilic character.

**Figure 1.**
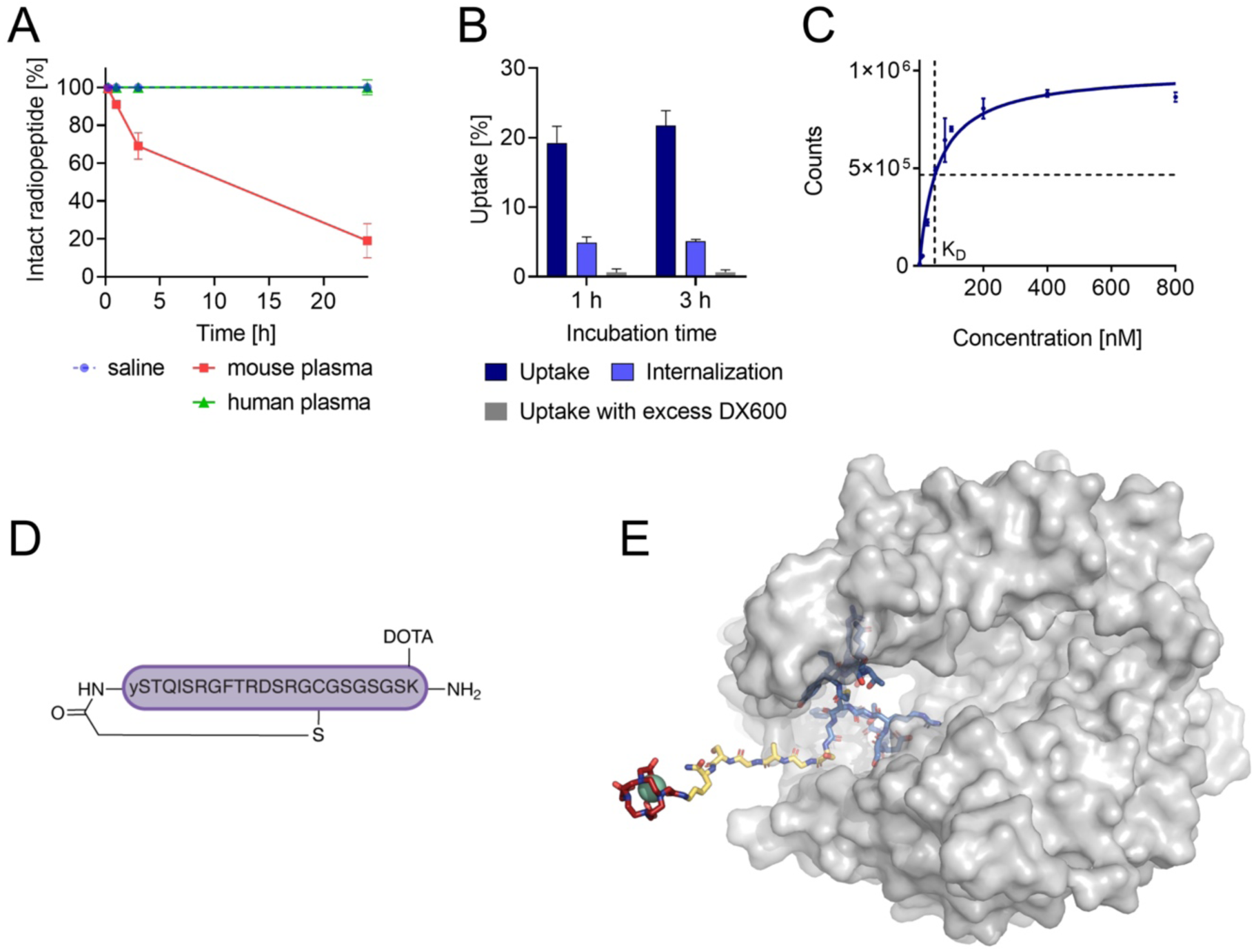
*In vitro* characterization and structural properties of [^67^Ga]Ga-DOTA-WJL-63. **A)** Stability of [^67^Ga]Ga-DOTA-WJL-63 in mouse and human blood plasma expressed as percent of intact radiopeptide relative to the values obtained in saline (set as 100%). **B)** Cell uptake and internalized fraction of [^67^Ga]Ga-DOTA-WJL-63 in HEK-ACE2 cells and blocked uptake of [^67^Ga]Ga-DOTA-WJL-63 in presence of excess DX600. **C)** Representative saturation binding curve of [^67^Ga]Ga-DOTA-WJL-63 on HEK-ACE2 cells (one representative example out of three replicates). **D)** Sequence of the cyclic peptide WJL-63. **E)** Molecular model of [^67^Ga]Ga-DOTA-WJL-63 bound to ACE2 (see Methods). The peptidase domain of ACE2 is shown as a transparent surface, WJL-63 (blue), the linker (yellow) and DOTA (red) are shown as sticks, and ^67^Ga is shown as a cyan sphere.

### Cell uptake and ACE2-binding affinity of the radiopeptide

In HEK-ACE2 cells the uptake of [^67^Ga]Ga-DOTA-WJL-63 reached 21 ± 2% and 23 ± 4% after an incubation period of 1 h and 3 h, respectively. The internalized fraction reached 4.9 ± 0.8% and 5.1 ± 0.3% after 1 h and 3 h, respectively, while the uptake was almost entirely prevented (<1%) by co-incubation of the HEK-ACE2 cells with excess DX600 (Figure 1B). The ACE2-binding affinity of [^67^Ga]Ga-DOTA-WJL-63 had a K_D_-value of 90 ± 28 nM (Figure 1C). The peptide sequence is shown in Figure 1D. A model of the WJL-63 peptide attached to DOTA via a linker (Figure 1E) shows that the addition of the DOTA chelator is not expected to interfere with WJL-63 binding to the protein.

### Structure of the ACE2 - WJL-63 peptide complex

To investigate the binding mode of WJL-63, we elucidated the crystal structure of the human ACE2 - WJL-63 peptide complex. For crystallization, the parent macrocyclic peptide, without the DOTA chelator, was used. The construct for crystallization comprised the extracellular part of human ACE2, including the peptidase domain and the noncatalytic domain, as in Towler et al.,^23^ and in addition contained a C-terminal His-tag to facilitate protein purification. The protein was expressed by secretion from insect cells and purified by Ni-NTA IMAC and size-exclusion chromatography (SEC). The resulting protein was crystallized by sitting drop vapor diffusion. A full dataset of a single crystal was recorded to a resolution of 2.2 Å at the Swiss Light Source (SLS) synchrotron at the PSI. The space group was C 1 2 1 and the structure was solved by molecular replacement with one copy of the complex in the asymmetric unit, followed by iterative steps of model building and refinement. Data collection and refinement statistics are shown in Supplementary Table 1.

### Overall structure and binding mode

The extracellular part of ACE2 comprises two domains, an N-terminal catalytic domain (residues 19 - 611) and a C-terminal non-catalytic collectrin homology domain (residues 612 - 740).^23^ The two subdomains of the catalytic domain flank the long substrate-binding cleft and are connected by an α-helical hinge. Our structure revealed that WJL-63 binds towards one side of the main peptide binding cleft of the peptidase domain of ACE2 in the open conformation (Figure 2).

**Figure 2.**
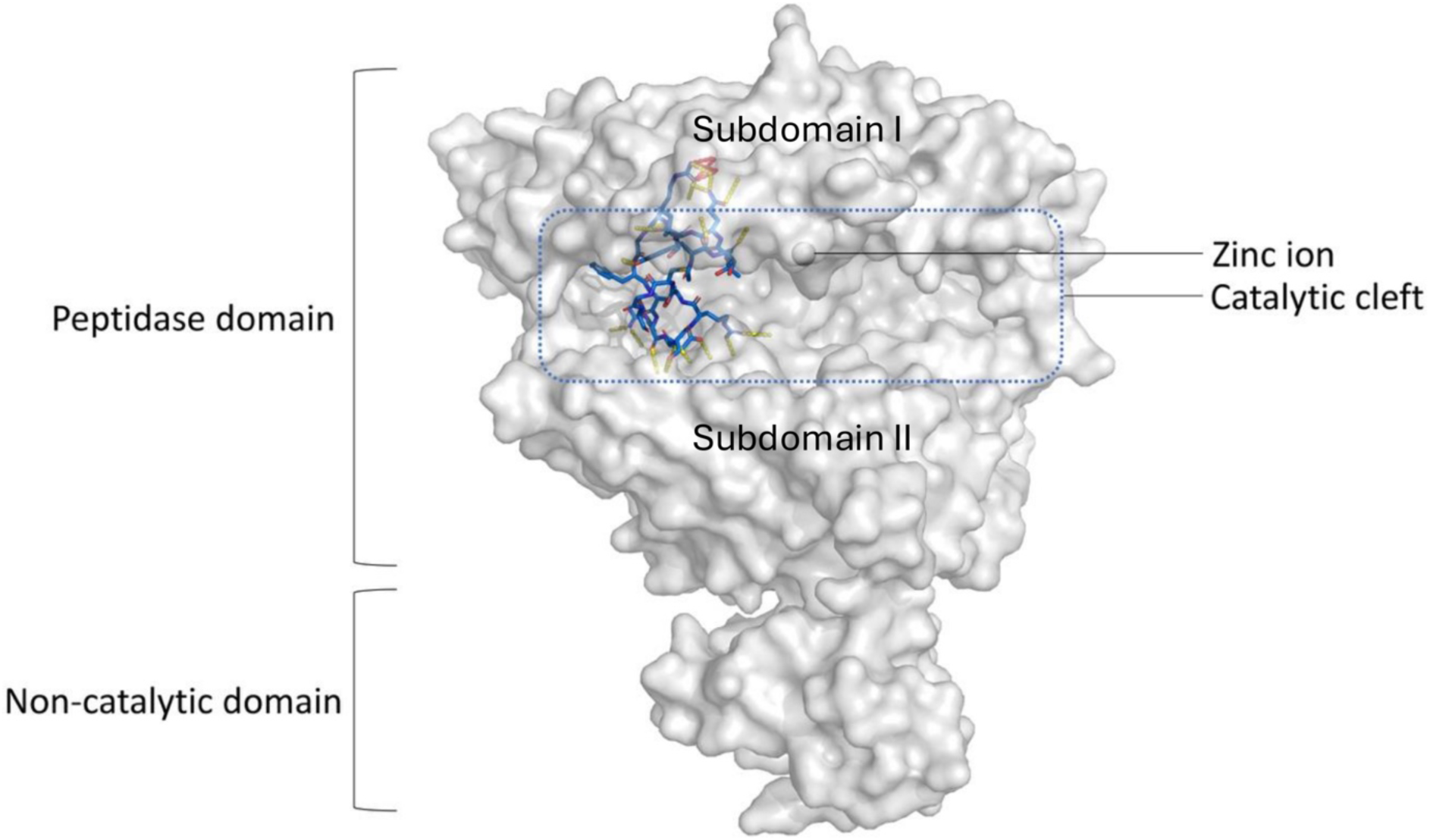
Crystal structure of the ACE2 - WJL-63 complex. ACE2 is shown as a semi-transparent surface representation in white. WJL-63 is shown as sticks with carbon atoms colored light blue, nitrogen atoms dark blue, oxygen atoms red and sulfur atoms yellow. Yellow dotted lines indicate hydrogen bonds between peptide and protein residues (see Supplementary Table 2 for details). The macrocyclic peptide binds to one side of the wide catalytic binding cleft in the open ACE2 conformation. The peptidase domain consists of two sub-domains that are connected by a hinge.

A surface area of 1142.5 Å^2^ is buried at the peptide-protein interface, according to analysis with PDBePISA.^24^ All peptide residues in the cyclic part of WJL-63 are clearly defined in the structure. The peptide interacts with sub-domain I of ACE2 through one edge, and with sub-domain II through the opposite edge. This binding mode requires an open ACE2 conformation.

The glycine-serine linker, which constitutes the non-cyclic part of the peptide, is only partially and weakly defined in the electron density. There is no overlap between the WJL-63 binding site and the binding site for the SARS-CoV-2 spike receptor-binding domain, which is located on top of sub-domain 1 of the peptidase domain^25, 26^ (Figure 3 A).

**Figure 3.**
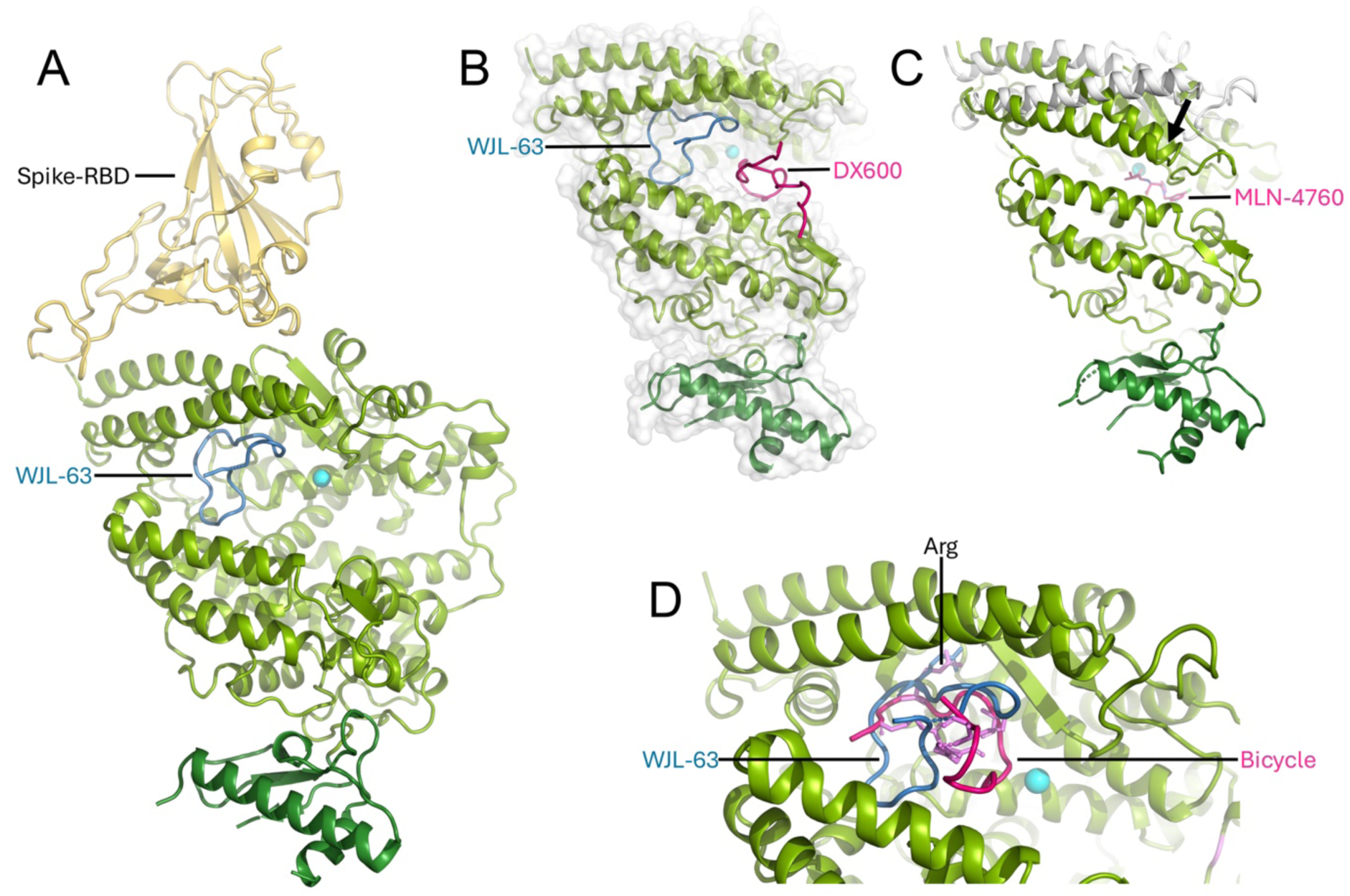
Comparison of the binding modes of WJL-63 and other ACE2-binders. **A)** The SARS-CoV-2 RBD binding site is located on the top of sub-domain 1 of the peptidase domain, which does not overlap with the catalytic cleft where WJL-63 binds. **B)** WJL-63 (blue) and DX600 (pink) binding sites. WJL-63 binds in the catalytic cleft of an open ACE2 conformation without contact to the conserved zinc binding site. The WJL-63 (blue) binding mode is distinct from the binding mode predicted for the peptide DX600 (pink) by modeling.^27^ The ACE2 - DX600 complex model suggests direct contact with the zinc binding site. **C)** MLN-4760 binds to the catalytic cleft in a closed ACE2 conformation, with direct contact to the zinc binding site.^23, 28^ The position of subdomain 1 in the open conformation (white cartoon) is shown for comparison. The downward movement in the closed state is indicated by the arrow. **D)** A bicyclic peptide (peptide shown as a cartoon in pink, scaffold shown as sticks in violet) recently identified by phage display^17^ (PDB:8BFW) shows overlap with WJL-63 (blue) in the binding mode, also binding to an open ACE2 conformation, but expanding more closely towards the catalytic zinc ion site. Arg 3 (shown as sticks in violet) of the bicyclic peptide interacts in the ACE2 binding pocket at a position that is equivalent to the interaction of WJL-63 Arg 7 (blue). The following structures were superposed onto the ACE2 – WJL-63 structure for the Figures: A) PDB:6LZG; B) model of ACE2 – DX600 structure (Ref.^27^); C) PDB:9FMM; D) PDB:8BFW. The zinc ion is shown in cyan. The catalytic domain of ACE2 is depicted in light green. The non-catalytic domain is shown in dark green. The structure in B is depicted as a cartoon with a semi-transparent surface. All other structures are shown as cartoons.

WJL-63 binds further towards the side of the catalytic cleft compared to the predicted binding of the peptide DX600,^27^ a peptide that was cyclized through a disulfide bond^16^ and was predicted to directly interact with the catalytic zinc ion (Figure 3B).

The binding mode of WJL-63 to the open ACE2 conformation is quite different from the binding mode of small molecule carboxypeptidase inhibitors such as MLN-4760, which stabilize the closed ACE2 conformation through interactions with cleft residues from both sub-domains^15, 23, 28^ (Figure 3C). We recently also solved a crystal structure of the extracellular domains of human ACE2 in complex with a fluorinated MLN-4760 derivative^28^ (PDB:9FMM). The overall binding mode of the fluorinated compound agrees with the binding mode previously described for MLN-4760,^23^ with direct contact to the conserved zinc ion site. Interestingly, in our structure, the zinc ion was present only with a partial occupancy, suggesting that the presence of zinc may not be required for compound binding. WJL-63 binds more towards one side of the ACE2 binding cleft and unlike MLN-4760 does not directly interact with the catalytic zinc ion site. WJL-63 hence only occupies a part of the catalytic cleft.

The binding mode of recently published bicyclic peptides (short linear peptides constrained with a chemical scaffold), which had been identified by phage display and optimized based on structural information^17^ (PDB:8BFW; PDB:8BYJ; PDB:8BN1; PDB:8B9P), show some overlap and similarities with our single-chain, macrocyclic WJL-63 (Figure 3D). Both, the bicyclic peptides and the macrocyclic peptide, bind to the open ACE2 conformation, and arginine 7, an important anchor for WJL-63 binding, has an equivalent (arginine 3) in the bicyclic peptides, supporting the critical importance of this residue for the binding to ACE2. Beyond this, the binding modes of the two peptides differ significantly, with WJL-63 being far more distant from the zinc ion site. Ser 7 of the bicyclic peptides in the structures containing a catalytic zinc ion (PDB:8BYJ; PDB:8BN1) approaches the zinc ion to a distance of 4.7 - 5.2 Å. The closest distances by which the macrocyclic peptide approaches the zinc ion in our ACE2 – WJL-63 complex structure are 11.7 Å (from WJL-63 Arg 14) and 12.1 Å (from Gln 4).

### Interaction details

The protein-peptide binding interface involves an extensive network of polar interactions (Figure 4A).

**Figure 4.**
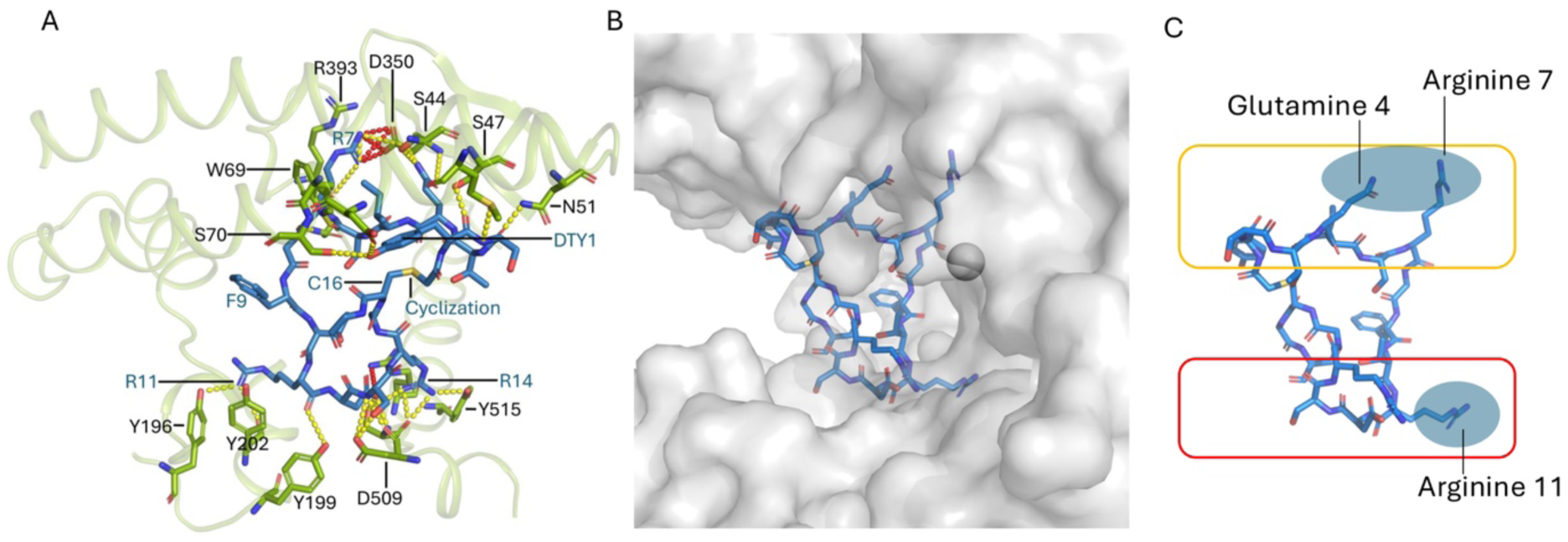
Interaction details of the ACE2 - WJL-63 complex. **A)** Hydrogen bonds (yellow dashed lines) and salt bridges (red dashed lines) (see Supplementary Table 2 for more details). **B, C)** Anchoring residues in the context of the ACE2 surface (B) and shown schematically (C), emphasizing interactions with the upper and lower sub-domains of the protein.

Hydrogen bonds and salt bridges formed between the peptide and ACE2 are listed in Supplementary Table 2. Supplementary Table 3 and 4 list the accessible surface area, buried surface area and solvation energy effect of WJL-63 residues and ACE2 residues, respectively, as determined using PDBePISA.^24^ Glutamine 4, arginine 7, arginine 11 (and arginine 14) of the peptide reach deep into the catalytic cleft (Figure 4 B and C).

## Discussion

We developed a macrocyclic peptide, WJL-63, that binds to ACE2 with relatively high affinity via a unique binding mode. The binding site was identified on one side of the catalytic cleft in the open ACE2 conformation, and the peptide did not directly contact the catalytic zinc ion site. This contrasts with other known ACE2 inhibitors, such as MLN-4760, which induces a closed enzyme conformation upon binding. Notably, the WJL-63 binding site does not overlap with the binding site of the SARS-CoV-2 spike receptor-binding domain, which is situated atop subdomain I of the ACE2 peptidase domain.

Bicyclic peptides with a nearby binding site to WJL-63 were recently described in the literature by Harman et al..^17^ They feature a peptide constrained into a bicyclic structure via a chemical scaffold. Similar to WJL-63, the bicyclic peptides bound to one side of the catalytic cleft in the open ACE2 conformation, sharing an anchoring interaction involving an arginine residue. However, the overall binding modes are different, with the bicyclic peptide extending closer to the catalytic zinc ion site, although the presence of a zinc ion was not required for binding of the bicyclic peptides, as some of the complex structures were solved in the absence of zinc.^17^

The rigid cyclic structure of WJL-63 appears to stabilize the open conformation of ACE2 through interactions with cleft residues from subdomains I and II. This leaves portions of the peptide accessible on the surface, making it well-suited for derivatization, for instance with a chelator for radiolabeling or a fluorophore. This was exemplified by derivatizing WJL-63 with a DOTA chelator for radiometal labeling, which did not prevent ACE2 binding to the peptide. It was further shown that co-incubation of the WJL-63 radiopeptide with DX600 prevents its uptake. This could be explained by slightly different ACE2 conformations required for binding of each of the peptides, as their binding sites are not expected to overlap. Overall, the DOTA-derivatized WJL-63 peptide showed promising *in vitro* characteristics after radiolabeling, especially regarding ACE2 selectivity and affinity, which was slightly higher than for the previously developed DX600-based radiopeptides.^27^ The WJL-based radiopeptide also exhibited more hydrophilic characteristics than DX600-based radiopeptides (logD ∼ −3 (Ref^27^) and overall high radiolytic stability. The new binding mode and higher affinity towards ACE2 of the radiopeptide may be advantageous when targeting physiological levels of the ACE2 enzyme which are commonly very low.

WJL-63 provides a strong basis for the development of novel reagents that target ACE2, for example fluorescent and/or electron-dense tags for *in situ* structural biology and multiscale imaging or radiopeptides for positron emission tomography (PET) imaging. Complementary to the bicyclic peptides reported in the literature,^17^ WJL-63 has potential to be used as a compound to study effects of ACE2 conformation by stabilizing a specific open enzyme state.

## Materials and Methods

### RaPID display

A library of ACE2-specific macrocyclic peptides was generated using the RaPID selection process in the laboratory of Prof. Hiroaki Suga at the University of Tokyo, Japan, by the standard protocol.^22^ Next generation sequencing (NGS) was applied for the enriched library at 3–6 rounds, and the sequences were aligned to identify families of potential ligand sequences. Among them, one of the most abundant families, WJL-63 was chosen for further evaluation. WJL-63 consists of <u>ClAc</u>-D-Tyr-Ser-Thr-Gln-Ile-Ser-Arg-Gly-Phe-Thr-Arg-Asp-Ser-Arg-Gly-<u>Cys</u>-Gly-Ser-Gly-Ser-Gly-Ser-Lys-NH_2_ where the N-terminus chloroacetyl group (underlined) reacted with the sulfhydryl group of the sidechain in the downstream cysteine residue (underlined) to selectively form a thioether bond.

### Peptide synthesis

WJL-63 was produced by SPPS using the following Fmoc-protected amino acids with side-chain protecting groups: Fmoc-Cys(Trt)-OH, Fmoc-Asp(O^t^Bu)-OH, Fmoc-Phe-OH, Fmoc-Gly-OH, Fmoc-Ile-OH, Fmoc-Lys(Boc)-OH, Fmoc-Gln(Trt)-OH, Fmoc-Arg(Pbf)-OH, Fmoc-Ser(^t^Bu)-OH, Fmoc-Thr(^t^Bu)-OH and Fmoc-D-Tyr(^t^Bu)-OH. Before starting the synthesis, Fmoc-Rink-Amide resin (0.56 mmol/g) was swollen in dimethylformamide (DMF) for 20 min. Fmoc deprotection was performed with 20% piperidine in DMF (2 x 10 min). Coupling was performed with Fmoc-protected amino acids (4.0 equiv to resin loading), *O*-(1*H*-6-Chlorobenzotriazole-1-yl)-1,1,3,3-tetramethyluronium hexafluorophosphate (HCTU, 3.9 equiv) as the coupling reagent and 4-methylmorpholine (NMM, 8.0 equiv) for 45 min. Capping was performed with 20% Ac_2_O and 10% NMM in DMF (10 min) after each coupling cycle. The peptide was synthesized with Symphony X after manual loading of the first amino acid, while chloroacetic acid coupling and peptide cyclization were performed manually. For the coupling of chloroacetic acid (4.0 equiv), oxyma (4.4 equiv) and diisopropylcarbodiimide (DIC, 4.0 equiv) were dissolved in a minimal amount of dimethylformamide and added to the resin. The reaction was purged with nitrogen gas for 45 min. Total deprotection and cleavage from the resin was achieved with a deprotection cocktail composed of 95% trifluoroacetic acid (TFA), 2.5% triisopropylsilane (TIPS) and 2.5% H_2_O (Milli-Q water, 15 mL of cleavage cocktail for 1 g of resin). The cleavage cocktail was concentrated under reduced pressure and the crude product was washed and precipitated three times with cold Et2O. The crude product was dissolved in H_2_O: ACN 1:1 with 0.1% TFA solution and purified with RP-HPLC. The Jasco HPLC system was equipped with a C18-MGII column (Capcell pak, 5 µm, 2.0 mm i.d. x 250 mm, Shiseido, Osaka, Japan) using a linear gradient of 5–75% acetonitrile (ACN) in H_2_O over 30 min at a flow rate of 10 mL/min.

The purified linear peptide (15 mg, 6.07 µmol) was dissolved in H_2_O (8 mL). Tris(2-carboxyethyl)phosphine (TCEP, 320 µL of 500 mM solution in water) and triethylamine (160 µL) were added and the pH of the solution was adjusted to 8 with additional triethylamine. The reaction was allowed to proceed at room temperature for 30 minutes. The reaction mixture was then acidified with TFA and further purified via RP-HPLC on a C18 column (5–75% ACN in H₂O).

The chelator DOTA-tris(tert-butyl ester) was conjugated to the cyclized peptide (1.0 equiv) using the chelator (2.0 equiv), HATU (1.9 equiv), and DIPEA (4.0 equiv). All reagents were dissolved in a minimal amount of N-methylpyrrolidone (NMP). The reaction was quenched after 2 hours at room temperature with a 1:1 mixture of H_2_O:ACN containing 0.1% TFA and purified via RP-HPLC on a C18 column (5–75% ACN in H_2_O). The protecting groups on DOTA were removed by treating the product with 95% TFA and 5% H_2_O for 2 hours at room temperature. The deprotection cocktail was then concentrated under reduced pressure. The remaining mixture was dissolved in a 1:1 solution of H₂O:ACN containing 0.1% TFA and purified again via RP-HPLC (5–75% ACN in H₂O). HRMS analysis confirmed the identity of the final product. The chemical structure of DOTA-WJL-63 is shown in Figure 5.

**Figure 5.**
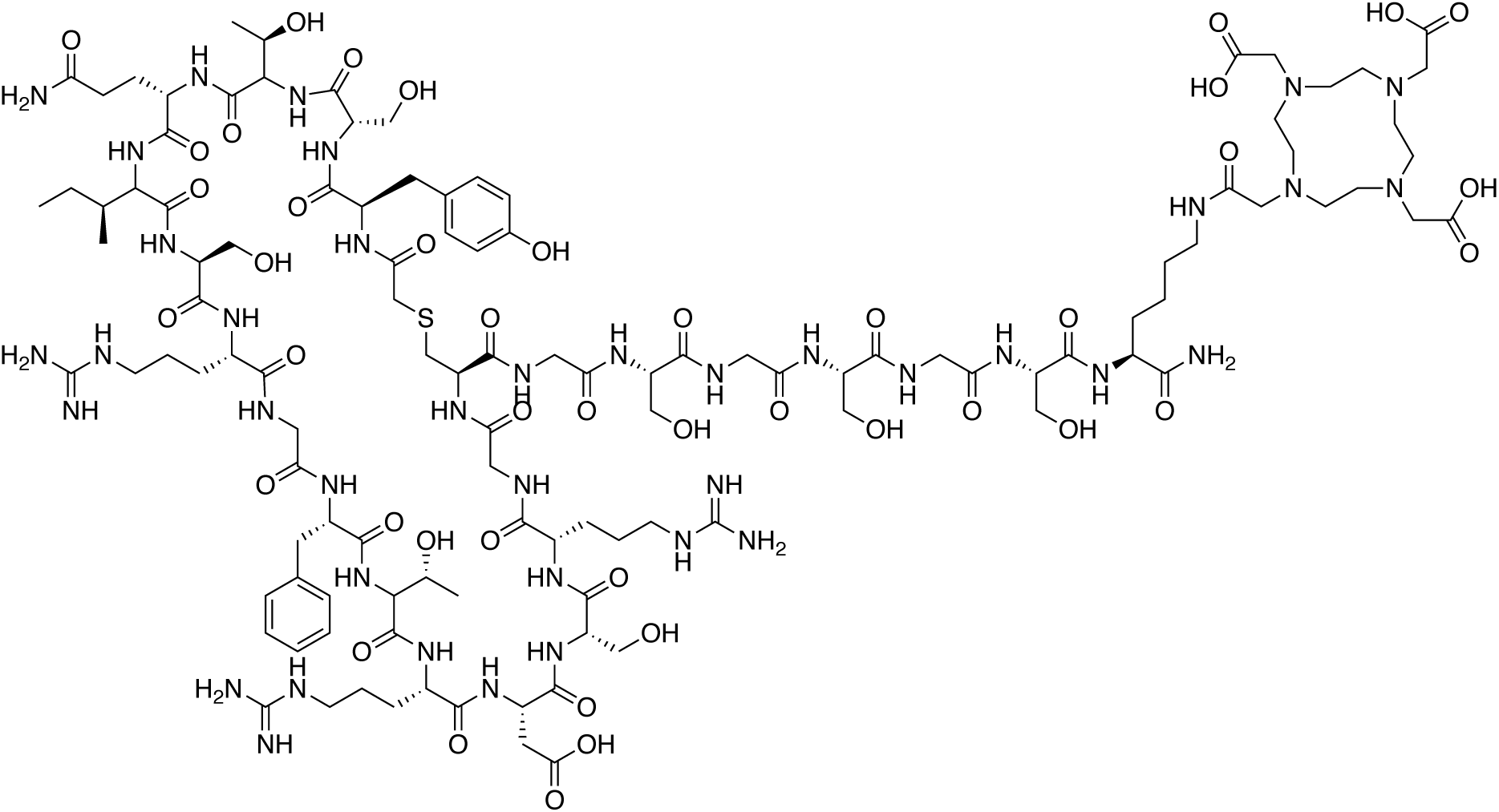
Chemical structure of DOTA-WJL-63 (D-Tyr-Ser-Thr-Gln-Ile-Ser-Arg-Gly-Phe-Thr-Arg-Asp-Ser-Arg-Gly-Cys-Gly-Ser-Gly-Ser-Gly-Ser-Lys-DOTA cyclized between the cysteine and the chloroacetic acid).

### Radiolabeling of the ACE2-targeting peptide

To enable radiolabeling of the peptide at high molar activity, the commercially obtained gallium-67 (Curium Netherlands B.V., the Netherlands, via b.e.imaging GmBH, Switzerland) was post-purified at the PSI according to a previously published procedure.^29^ The purified [^67^Ga]GaCl_3_ in HCl (∼0.1 M) was added to a sodium acetate solution (0.5 M, pH 8) to obtain a buffered solution with a pH of ∼4. After addition of DOTA-WJL-63 to obtain a molar activity of 5-20 MBq/nmol, the reaction mixture was incubated at 95 °C for 10 min. Quality control of the radiolabeled WJL-63 peptide was performed after dilution of the radiolabeling mixture in Milli-Q water containing sodium diethylenetriamine pentaacetic acid (Na_5_-DTPA; 50 μM) on a Merck Hitachi LaChrom HPLC system equipped with a D-7000 interface, a L-7200 autosampler, a radioactivity detector (LB 506 B; Berthold) and a L-7100 pump connected with a C18 column (Xterra, 5 μm, 4.6×150 mm, Waters, Milford, MA, USA). The radiopeptide was eluted using a linear gradient of Milli-Q water containing 0.1% TFA (95–20%) and ACN (5–80%) over 15 min at a flow rate of 1.0 mL/min. The radiopeptide was used directly for *in vitro* studies without further purification steps.

### Stability testing of DOTA-WJL-63

The stability of [^67^Ga]Ga-DOTA-WJL-63 (20 MBq/nmol) formulated in saline, was tested *in vitro* over 24 h at room temperature at a concentration of 10 MBq/100 µL formulated in saline, with and without the presence of the radical scavenger ascorbic acid using the HPLC system described above. The HPLC chromatograms were analyzed by determination of the peak area of the radiolabeled peptide, the released gallium-67, as well as degradation products of unknown structure. The quantity of the intact product was expressed as the percentage of the sum of integrated peak areas of the entire chromatogram and set in relation to the original value determined at t = 0, which was set as 100%.

Additionally, the integrity of [^67^Ga]Ga-DOTA-WJL-63 (20 MBq/nmol) in mouse and human blood plasma (Rockland Immunochemicals, Inc. PA, USA and Blood donation SRK Aargau-Solothurn, Switzerland) was determined using thin layer chromatography (TLC). Reversed phase C-18 plates (TLC Silica gel 60 RP-18; Merck) were used as the stationary phase and a mixture of citrate buffer (pH 5.5; 0.1 M) and acetonitrile (7:3; *v/v*) as the mobile phase. Under these conditions, uncoordinated gallium-67 migrated with the front while the radiopeptide and potential larger fragments migrated to the center of the plate. After radiolabeling, the radiopeptide was incubated in saline, mouse or human blood plasma (10 MBq/200 µL), in the presence of ascorbic acid to avoid radiolytic degradation at 37 °C. TLC was performed with an aliquot of the control sample in saline and the plasma samples after 15 min, 1 h, 3 h and 24 h. The TLC plates were exposed to a phosphor screen (super resolution screen PSR10450013) followed by development using a storage phosphor system (Cyclone Plus, PerkinElmer). The quantification of the signals was carried out using the OptiQuant software (version 5.0, Bright Instrument Co Ltd., PerkinElmer). The obtained chromatograms were analyzed by determination of the peak area of the radiolabeled peptide, the released gallium-67 as well as degradation products of unknown structure. The quantity of the intact product was expressed as the percentage of the sum of integrated peak areas of the entire chromatogram and set in relation to the value determined at the respective timepoints of the radiopeptide diluted in saline which was set as 100% to correct for TLC artefacts. The results were then expressed as average ± SD from n = 3 independent experiments.

### Determination of the distribution coefficient (logD)

The distribution coefficient (logD value) of [^67^Ga]Ga-DOTA-WJL-63 was determined using a shake-flask method as previously reported for DX600 radiopeptides.^27^ The distribution coefficients were indicated as the average of n = 3 independent measurements ± standard deviation (SD), each performed with five replicates.

### Cell culture

HEK-293 cells transfected with ACE2, referred to as HEK-ACE2, were obtained from Innoprot (Innovative Technologies in Biological Systems S.L. Bizkaia, Spain). The cells were cultured in Dulbecco’s Modified Eagle Medium (DMEM) supplemented with non-essential amino acids, fetal calf serum and antibiotics. Hygromycin B was used to maintain the expression of ACE2 which was previously verified^27^ by western blotting. These HEK-ACE2 cells were cultured under standard conditions at 37 °C and 5% CO_2_ and sub-cultured using a phosphate buffered saline (PBS)/ethylenediaminetetraacetic acid (EDTA) and trypsin mixture to release cells from the culture flasks.

### Cell uptake studies

The uptake and internalization of [^67^Ga]Ga-DOTA-WJL-63 was determined using a previously published protocol.^27^ In brief, HEK-ACE2 cells were seeded in 12-well-plates (1 × 10^6^ cells in 2 mL DMEM medium with supplements/well) allowing cell adhesion and growth overnight. The cells were rinsed with PBS prior to the addition of DMEM medium without supplements (975 µL/well) and [^67^Ga]Ga-DOTA-WJL-63 (20 MBq/nmol) in a volume of 25 µL (1.9 pmol, 38 kBq). Cells in half of the wells were co-incubated with excess DX600 (2 µM, Selleckchem), which was found to prevent WJL-63-binding, to block ACE2 on the cell surface. After incubation of the cells for 1 h or 3 h at 37 °C and 5% CO_2_, the cells were rinsed with cold PBS to determine total uptake of the radiopeptide. To assess the internalized fraction, a stripping buffer (acidic glycine buffer containing saline, pH 2.8) was applied to release ACE2-bound radiopeptide from the cell surface. Cell samples were lysed using NaOH (1 M, 1 mL) and measured in a γ-counter (Wallac Wizard 1480, PerkinElmer). The concentration of proteins in each well was determined using a Micro BCA Protein Assay kit (Pierce, Thermo Scientific). The results were expressed as percentage of total added activity and each well was standardized to the average content of protein per well of each plate.

### Determination of ACE2-binding affinity

The K_D_ values indicating the ACE2-binding affinity of [^67^Ga]Ga-DOTA-WJL-63 was determined using HEK-ACE2 cells cultured overnight in 48-well-plates (0.25 x 10^6^ cells per well, in 500 µL DMEM medium with supplements). During the entire experimental procedure performed the next day, the well-plates were kept on ice. The cells were rinsed with ice-cold PBS before addition of various concentrations of the radiolabeled peptide (5 MBq/nmol; 1–2000 nM peptide) in ice-cold DMEM medium without supplements. Half of the cell samples were co-incubated with DX600 (20 μM, 20 nmol) to block ACE2, enabling the determination of non-specific binding. After incubation of the well-plates for 1 h at 4 °C, the cells were rinsed twice with ice-cold PBS, lysed using NaOH (1 M, 600 µL) and counted for activity in a γ-counter (Wallac Wizard 1480, PerkinElmer). The K_D_ values were determined by plotting specific binding (total binding minus unspecific binding determined with blocking agent) against the molar concentration of the added radiopeptide. The nonlinear regression analysis was performed using GraphPad Prism software (version 8.0) and the results were expressed as average ± SD from K_D_-values obtained from n = 3 independent experiments.

### ACE2 construct design

The construct for crystallization comprised the extracellular part of human ACE2, including the peptidase domain and the noncatalytic domain as in^23^ and in addition contained a C-terminal His-tag to facilitate protein purification. A synthetic gene, codon optimized for expression in insect cells, coding for amino acids 1-740 of human ACE2 (UniProt sp Q9BYF1), including the signal sequence and a non-cleavable C-terminal 8xHis-tag, was purchased from Genewiz/Azenta Life Sciences, cloned into the NotI and KpnI sites of the vector pJM-24_pAC8Red-eGFP-His8.^30^

### Protein expression and purification

SF9 cells were co-transfected with the plasmid coding for the ACE2 construct and linearized Bac 10 KO1629 DNA in SF900 II medium for virus generation. After viral amplification, 0.2 µM filtered V2 viral stock was used to infect Hi5 cells for protein expression.

The protein was expressed by secretion from baculovirus-infected Hi5 insect cells in SF900 II medium at 27 °C in a shakeflask. The cleared supernatant was loaded onto a 5 ml HisTrap Excel column (Cytiva) equilibrated in 5 column volumes of 50 mM TRIS pH 7.4, 500 mM NaCl and washed with 20 column volumes of 50 mM TRIS pH 7.4, 500 mM NaCl, 10 mM imidazole, at 4 °C. Elution was carried out using 5 column volumes of 50 mM TRIS pH 7.4, 500 mM NaCl, 500 mM imidazole. The eluted protein was concentrated in a 50 kDa cutoff concentrator (Millipore) and further purified over a Superdex 200 increase 10/300 column in 20 mM TRIS pH 7.4, 150 mM NaCl. Glycerol was added to a final concentration of 10% and aliquots were flash-frozen in liquid nitrogen. Prior to crystallization trials, aliquots were thawed on ice, centrifuged at 13’000 rcf and purified over a Superose 6 increase 10/300 column in 20 mM TRIS pH 7.4, 150 mM NaCl. The protein was then concentrated to 8.5 mg/ml in a 50 kDa cutoff concentrator (Millipore). ZnCl_2_ was added to a final concentration of 1 mM.

### Crystallization, data collection and structure elucidation

The protein-peptide complex was formed by addition of a 1.2x molar excess of WJL-63 to SEC purified, concentrated (8.5 mg/ml) human ACE2 protein, followed by incubation on ice for 30 minutes. The protein was crystallized by sitting drop vapor diffusion using a Mosquito robot. The drop size was 100 nl protein plus 100 nl reservoir solution. Optimal crystals formed in well E9 of the Qiagen PEGs II Suite crystal screen (0.2 M ammonium sulfate, 0.1 M HEPES pH 7.5, 16% (w/v) PEG 4000, 10% (v/v) isopropanol). The cryo buffer consisted of reservoir solution plus 40% ethylene glycol. A complete dataset from a single crystal was recorded at 100 K to a resolution of 2.2 Å at the X06sa beamline of the Swiss Light Source (SLS) synchrotron at PSI, Villigen PSI, Switzerland, using a wavelength of 1.000 Å. Data processing was carried out using XDS.^31^ The structure was solved by molecular replacement with Phaser^32^ using an AlphaFold model^33, 34^ of human ACE2 as the search model. The space group was C 1 2 1. The asymmetric unit contained one copy of the complex. Regions of the initial model that did not fit the electron density were deleted, and after additional refinement, the missing parts were modeled in Coot.^35^ In subsequent refinement cycles, density corresponding to the WJL-63 macrocyclic peptide became apparent, which allowed manual building of the peptide. The model was then optimized through iterative steps of model building in Coot and refinement using phenix.refine.^36^ MolProbity^37^ was used to validate the geometry and stereochemistry. Structure figures were prepared using PyMOL (The PyMOL Molecular Graphics System, Version 3.1.0 Schrödinger, LLC) and Coot. The binding interface was analyzed using PDBePISA^24^ v1.52.

### Molecular modeling

The molecular model of DOTA-WJL-63 was built using the crystal structure of the WJL-63 peptide bound to ACE2 reported in this work as a template. We used molecular graphics software (The PyMOL Molecular Graphics System, Version 3.1.0, Schrödinger, LLC; Avogadro, Version 1.2.0) to model the glycine-serine linker (SGSGSK) and the DOTA chelator. The geometry of these components was optimized using the MMFF94 force field. This optimization process involved iteratively adjusting atomic positions to minimize steric clashes and ensure chemically realistic bond lengths and angles.

## Acknowledgements

We thank the MX group for support at the Swiss Light Source (SLS) beamlines at PSI and Jonas Mühle at the PSI for advice and support with insect cell cultures. The authors also thank Fan Sozzi-Guo and Susan Cohrs for technical support during the preclinical evaluation of the radiopeptide as well as Nicholas P. van der Meulen and Pascal V. Grundler for establishing the gallium-67 post-purification method. This work was supported by grants from the Promedica Stiftung Chur (1401/M), and the Swiss National Science Foundation (SNF SPARK, CRSK-3_190414) to R.M.B.. D.B. was also funded by the Swiss National Science Foundation NRP78 program (Grant No. 310030_188978: PI: C.M.). The project was furthermore supported by the Japan Society for the Promotion of Science (JSPS) Grant-in-Aid for Specially Promoted Research (JP20H05618 to H.S.).

## Author contributions

R.M.B. and M.W. produced, purified and crystallized the protein-peptide complex. RMB collected the X-ray diffraction data and solved the structure. R.M.B. and M.J.R. refined the structure. A.A. and H.S. carried out the RaPID selection campaign to select and identify the macrocyclic peptide. J.W. and J.W.B. synthesized the macrocyclic peptide and the DOTA conjugate. D.B. and C.M. carried out the radiolabeling followed by the characterization of the radiopeptide and cellular experiments. X.D. modeled the DOTA-WJL-63. All authors contributed to writing and revising the manuscript led by R.M.B..

