## Supporting Information for "Development and structure-guided characterization of a novel ACE2-binding macrocyclic peptide"

**Supplementary Table 1) Crystallographic data collection and refinement statistics**

| | |
|---|---|
| Space group | C 1 2 1 |
| Cell dimensions | |
| *a, b, c* (Å) | 103.1, 89.1, 112.7 |
| $\alpha, \beta, \gamma$ (°) | 90.0, 108.3, 90.0 |
| Resolution range (Å) | 46.64 - 2.20 (2.279 - 2.20) |
| $R_{meas}$ | 0.5112 (2.07) |
| $CC_{1/2}$ | 0.98 / 0.57 |
| I/σI | 11.52 (1.78) |
| Completeness (%) | 98.75 (98.69) |
| Multiplicity | 6.9 (6.9) |
| Total reflections | 336921 (33480) |
| Unique reflections | 48618 (4822) |
| $R_{work}$ / $R_{free}$ | 0.1865 / 0.2257 |
| No. atoms | |
| Macromolecules | 5666 |
| Ligands | 357 |
| Solvent | 128 |
| B-factors (Å$^2$) | |
| Macromolecules | 53.81 |
| Ligands | 71.23 |
| Solvent | 49.72 |
| R.m.s. deviations | |
| Bond lengths (Å) | 0.011 |
| Bond angles (°) | 1.09 |

Highest-resolution shell in parentheses.

**Supplementary Table 2) Hydrogen bonds and salt bridges between WJL-63 and ACE2** (analyzed using PDBePISA.[24])

### Hydrogen bonds

| WJL-63 residue | Distance (Å) | ACE2 residue |
|---|---|---|
| DTY 1 (OH) | 2.77 | Asp 67 (OD1) |
| DTY 1 (OH) | 3.30 | Ser 70 (OG) |
| Ser 2 (N) | 3.28 | Met 62 (SD) |
| Gln 4 (NE2) | 2.68 | Asp 350 (O) |
| Gln 4 (NE2) | 3.01 | Ser 44 (OG) |
| Arg 7 (NH1) | 3.73 | Arg 393 (O) |
| Arg 7 (NH1) | 2.73 | Asp 350 (OD2) |
| Arg 7 (NH2) | 3.84 | Asp 350 (O) |
| Arg 7 (NH2) | 2.93 | Asp 350 (OD1) |
| Arg 11 (NH1) | 3.04 | Tyr 196 (OH) |
| Arg 11 (NH1) | 3.13 | Tyr 202 (0) |
| Ser 13 (N) | 3.40 | Asp 509 (OD2) |
| Ser 13 (N) | 2.86 | Asp 509 (O) |
| Ser 13 (OG) | 2.61 | Asp 509 (OD2) |
| Arg 14 (NH1) | 2.85 | Ser 511 (O) |
| Arg 14 (NH1) | 3.28 | Ser 511 (OG) |
| Arg 14 (NH2) | 3.03 | Tyr 515 (OH) |
| Arg 14 (NH2) | 2.98 | Ser 511 (O) |
| DTY 1 (O) | 2.27 | Ser 47 (OG) |
| Ser 2 (O) | 2.47 | Asn 51 (ND2) |
| Gln 4 (OE1) | 3.81 | Asp 350 (N) |
| Ser 6 (O) | 3.70 | Trp 69 (NE1) |
| Arg 7 (O) | 3.12 | Asn 394 (ND2) |
| Arg 11 (O) | 3.25 | Tyr 199 (OH) |
| Asp 12 (OD2) | 2.76 | Ser 511 (N) |
| Asp 12 (OD2) | 3.15 | Ser 511 (OG) |

### Salt bridges

| WJL-63 residue | Distance (Å) | ACE2 residue |
|---|---|---|
| Arg 7 (NH1) | 3.76 | Asp 350 (OD1) |
| Arg 7 (NH1) | 2.73 | Asp 350 (OD2) |
| Arg 7 (NH2) | 2.93 | Asp 350 (OD1) |
| Arg 7 (NH2) | 3.38 | Asp 350 (OD2) |
| Asp 12 (OD2) | 3.99 | Arg 514 (NH2) |

**Supplementary Table 3) Interfacing residues of the macrocyclic peptide** (analyzed using PDBePISA.[24])

| WJL-63 residue | Accessible Surface Area (ASA), Å$^2$ | Buried Surface Area (BSA), Å$^2$ | Solvation Energy Effect $\Delta^i G$ (kcal/mol) | Residue engaged in polar interaction |
|---|---|---|---|---|
| DTY 1 | 171.34 | 142.71 | 1.03 | H-bond |
| Ser 2 | 124.10 | 77.27 | 0.39 | H-bond |
| Thr 3 | 86.52 | 0.50 | 0.01 | --- |
| Gln 4 | 165.95 | 121.25 | -0.51 | H-bond |
| Ile 5 | 110.42 | 104.96 | 1.52 | --- |
| Ser 6 | 57.43 | 18.83 | -0.08 | H-bond |
| Arg 7 | 252.70 | 235.72 | -1.31 | H-bond / salt bridge |
| Gly 8 | 51.42 | 38.11 | 0.33 | --- |
| Phe 9 | 211.89 | 138.23 | 1.99 | --- |
| Thr 10 | 56.06 | 4.82 | -0.03 | --- |
| Arg 11 | 254.82 | 197.24 | -0.92 | H-bond |
| Asp 12 | 75.62 | 62.22 | 0.58 | H-bond / salt bridge |
| Ser 13 | 109.91 | 71.64 | -0.04 | H-bond |
| Arg 14 | 207.24 | 149.35 | -0.60 | H-bond |
| Cys 16 | 31.90 | 7.88 | -0.09 | --- |
| Gly 17 | 119.21 | 8.20 | 0.13 | --- |

**Supplementary Table 4) Interfacing residues of ACE2** (analyzed using PDBePISA.[24])

| ACE2 residue | Accessible Surface Area (ASA), Å$^2$ | Buried Surface Area (BSA), Å$^2$ | Solvation Energy Effect $\Delta^i G$ (kcal/mol) | Residue engaged in polar interaction |
|---|---|---|---|---|
| Phe 40 | 71.39 | 65.91 | 1.05 | --- |
| Ser 43 | 17.12 | 17.12 | -0.09 | --- |
| Ser 44 | 13.99 | 13.99 | 0.03 | H-bond |
| Ser 47 | 23.69 | 23.45 | -0.02 | H-bond |
| Tyr 50 | 48.26 | 11.57 | 0.19 | --- |
| Asn 51 | 28.74 | 22.25 | -0.29 | H-bond |
| Met 62 | 40.45 | 29.78 | 0.72 | H-bond |
| Asn 63 | 74.40 | 13.01 | -0.02 | --- |
| Gly 66 | 21.32 | 21.32 | 0.32 | --- |
| Asp 67 | 97.09 | 18.41 | -0.21 | H-bond |
| Trp 69 | 40.49 | 35.28 | 0.25 | H-bond |
| Ser 70 | 67.80 | 32.41 | -0.17 | H-bond |
| Leu 73 | 56.16 | 48.46 | 0.77 | --- |
| Lys 74 | 127.92 | 7.79 | 0.11 | --- |
| Ser 77 | 16.02 | 8.70 | 0.14 | --- |
| Ala 99 | 40.52 | 16.03 | -0.08 | --- |
| Leu 100 | 6.74 | 5.74 | 0.06 | --- |
| Gln 102 | 116.58 | 80.20 | -0.66 | --- |
| Leu 120 | 37.51 | 10.22 | 0.16 | --- |
| Lys 187 | 14.35 | 3.46 | -0.13 | --- |
| Tyr 196 | 58.18 | 15.46 | -0.18 | H-bond |
| Tyr 199 | 8.74 | 4.88 | -0.04 | H-bond |
| Tyr 202 | 37.10 | 26.45 | 0.22 | H-bond |
| Trp 203 | 36.37 | 25.82 | 0.12 | --- |
| Gly 205 | 19.99 | 14.17 | 0.01 | --- |
| Asp 206 | 63.67 | 16.93 | 0.19 | --- |
| Val 343 | 57.07 | 0.84 | 0.01 | --- |
| Ala 348 | 30.43 | 3.44 | -0.04 | --- |
| Trp 349 | 44.07 | 36.11 | 0.58 | --- |
| Asp 350 | 34.01 | 23.19 | -0.14 | H-bond / salt bridge |
| Gly 352 | 7.13 | 3.81 | 0.05 | --- |
| Tyr 385 | 7.84 | 0.61 | -0.01 | --- |
| Phe 390 | 36.07 | 24.88 | 0.28 | --- |
| Leu 391 | 22.89 | 20.20 | 0.31 | --- |
| Arg 393 | 42.33 | 23.56 | -0.05 | H-bond |
| Asn 394 | 69.12 | 28.06 | -0.36 | H-bond |
| His 505 | 16.41 | 0.12 | 0.00 | --- |
| Asn 508 | 28.18 | 7.74 | -0.09 | --- |
| Asp 509 | 27.17 | 25.13 | -0.07 | H-bond |
| Tyr 510 | 111.93 | 61.88 | 0.77 | --- |
| Ser 511 | 16.58 | 12.23 | -0.10 | H-bond |
| Arg 514 | 91.22 | 38.51 | -0.86 | Salt bridge |
| Tyr 515 | 52.63 | 6.91 | -0.08 | H-bond |
